# Cyclin B phosphorylation by PKA safeguards the G2-arrest in vertebrate oocytes

**DOI:** 10.64898/2026.08.03.742451

**Authors:** Martina Santoni, Tran Le, Veronique Legros, Guillaume Chevreux, Catherine Jessus, Enrico Maria Daldello

## Abstract

Entry into meiotic M-phase in vertebrate oocytes requires activation of Cdk1-Cyclin B, which is restrained by the cAMP-PKA signalling during prophase arrest. Although Cdc25C and ARPP19 are established PKA substrates in this context, they do not fully explain how PKA represses Cdk1 activation. Here we used Turbo-ID approach using catalytic PKA (PKAc) as the bait to identify substrates in *Xenopus laevis* oocytes. Unexpectedly, this screen identified Cyclin B2, the regulatory subunit of Cdk1, as a PKA-proximal protein. Although Cyclin B2 does not stably associate with PKAc, it harbors a conserved PKA motif around Serine 271 within Cyclin Box 2. We show that PKAc phosphorylates Cyclin B2 at S271 both *in vivo* and *in vitro*. Functionally, a phosphomimic Cyclin B2 mutant at this site fails to induce meiotic maturation and Cdk1 activation. S271 phosphorylation neither alters Cyclin B2 stability nor its binding to Cdk1 *in vivo*. S271 phosphorylation does not impair Cdk1 activity toward a single-site substrate, such as PP1, but slows Cyclin-B-dependent multisite phosphorylation of Cdc25C, a key regulator of the Cdk1 activation network. These findings identify Cyclin B as a direct and conserved PKA target, revealing the mechanistic link between PKA activity and Cdk1-Cyclin B repression that maintains oocyte prophase arrest.

## INTRODUCTION

In metazoans, oocytes undergo a prolonged arrest at prophase I of meiosis, ranging from a few weeks to several decades depending on the species. This extended period provides the opportunity for oocyte growth and for the accumulation of the nutritional reserves and maternal components, mRNAs and proteins, required to sustain early embryonic development following fertilization. This prophase I arrest, which corresponds to an extended G2 phase of the cell cycle, is therefore essential for the survival of animal species. Elucidating the molecular mechanisms that underlie its maintenance provides valuable insights not only into the regulation of sexual reproduction but also into the fundamental mechanisms governing cell cycle progression.

In vertebrate oocytes, prophase I arrest is maintained by cAMP-dependent Protein Kinase A (PKA), which indirectly represses activation of the Cdk1-Cyclin B complex, the master regulator of M-phase^1,2^. At the time of ovulation, a hormonal stimulus, progesterone in *Xenopus*, one of the most widely used model organisms for studying this process, lowers the level of cAMP, leading to PKA inhibition that allows meiosis to resume^3,4^. Experimentally, manipulating PKA activity is sufficient to control the decision of the cell to enter meiotic division. Overexpression of the PKA regulatory subunit (PKAr)^5^ or microinjection of PKI^6^, a specific inhibitor of PKA, promotes meiosis resumption, whereas increasing PKA catalytic subunit (PKAc) prevents it^5^. Although progesterone induces PKA inhibition within approximately 30 minutes, Cdk1 activation does not occur until 3–5 hours later. During this interval, the decrease in PKA activity initiates a program of *de novo* protein synthesis involving proteins such as cyclin B1, the kinase Mos, and Ringo, as well as additional, as yet unidentified, factors^7^. This translation program is essential for Cdk1 activation, which is a two-step process^8^. First, a “starter” amount of active Cdk1 is generated in a protein translation dependent manner; second, once this threshold is reached, a series of positive feedback loops drives the full activation of Cdk1, a process known as the Cdk1 auto-amplification loop^9^.

Once active, Cdk1-Cyclin B and its effector kinases efficiently phosphorylate a large set of substrates to execute M-phase entry^10–13^. Beyond the Cdk consensus motif (S/T-P-x-K/R)^14,15^, substrate specificity relies on docking modules on Cyclins and accessory factors as CKS^16^. CKS proteins bind phosphorylated serine (Ser) or threonine (Thr) residues to promote processive multi-site phosphorylation^17^, while Cyclin subunits contribute to substrate-class specificity^18–21^. A conserved MRAIL-containing hydrophobic patch on multe types of cyclins recognizes R-x-L docking motifs in substrates, positioning them for efficient phosphorylation by Cdk1^18,19,21,22^. Beyond direct substrate recruitment, this hydrophobic patch also directs Cyclin B–Cdk1 complexes to centrosomes, thereby contributing to the spatial organization of mitotic substrate phosphorylation^23^. Recent work uncovered a positively charged phosphate-binding pocket (PBP) on Cyclin B that binds phospho-Ser/Thr on substrates, enabling Cdk1 to phosphorylate additional non-consensus sites^24,25^.

PKA substrates repress multiple nodes of the protein network controlling Cdk1 activation. The phosphatase Cdc25C was the first physiological PKA target identified in *Xenopus* oocytes^26^. PKA phosphorylation of Cdc25C at Ser-287 creates a 14-3-3 binding site that functionally inactivates and sequesters Cdc25C, supporting prophase arrest. In mouse oocytes, Cdc25B is likewise restrained by PKA and 14-3-3-dependent mechanisms before Cdk1 activation^27^. Together these data place Cdc25C under PKA control. However, in *Xenopus*, the major S287 dephosphorylation occurs only after Cdk1 activation and is blocked by Cdk1 inhibition, indicating that Cdc25C dephosphorylation is involved in the activation of the Cdk1 auto-amplification loop but does not contribute to the generation of the initial threshold level of Cdk1 activity^26,28,29^. In mouse oocytes, Wee1B, an oocyte-specific Wee1 family kinase, acts downstream of PKA to maintain the prophase arrest. Ser-15 of Wee1B is a major *in vitro* PKA phosphorylation site, and phosphorylation at this residue enhances Wee1B inhibitory activity toward Cdk1^30^. Notably, in *Xenopus*, the oocyte-specific Wee1 family kinase, named Wee1A, accumulates only after metaphase I^31^. In *Xenopus* oocytes, PKA also phosphorylates Ringo, a direct activator of Cdk1^32,33^. PKA-mediated phosphorylation of Ringo promotes its degradation during prophase arrest, thereby preventing its accumulation and premature activation of Cdk1^34^. More recently, a fourth PKA substrate was identified, ARPP19^28^. PKA phosphorylation at S109 is necessary and sufficient to keep *Xenopus* oocytes in prophase^28^. Following progesterone stimulation, ARPP19 is partially dephosphorylated before Cdk1 activation, thereby permitting the “starter” step, while 3 to 5 hours later, Greatwall-dependent phosphorylation of ARPP19 at S67 initiates the auto-amplification loop independently of PKA^28,35^.

These established substrates, however, do not fully account for two key observations. First, downregulating PKA still initiates the protein-accumulation program upstream of Cdk1 activation, even when ARPP19 is held in a phospho-mimetic S109D state^29^. This observation implies that additional PKA effectors regulate translation/stability prior to Cdk1 activation. Second, overexpression of Cyclin B induces Cdk1 activation in the absence of additional protein synthesis, yet this activation remains sensitive to PKA inhibition^36^. This indicates that PKA inhibits Cdk1 activation also at steps downstream of the activation of protein translation. Thus, the full set of PKA targets that restrain Cdk1 activation, both upstream and at the level of the Cdk1 activation network, remains incompletely defined.

Here, we performed a Turbo-ID screening to identify novel PKA effectors during the prophase arrest in *Xenopus* oocytes. Unexpectedly, we uncovered Cyclin B2 as a putative PKA substrate. We demonstrate that Cyclin B2 is phosphorylated by PKA *in vivo* and *in vitro* at a highly conserved site across eukaryotes, Ser271 (S271). This phosphorylation suppresses the ability of Cyclin B2 to induce Cdk1 activation by preventing Cdk1 from phosphorylating key regulators within the kinase–phosphatase network that drives its own activation. These findings place Cyclin B2, alongside Cdc25C and ARPP19, as a direct PKA target and provide the first mechanistic link between PKA downregulation and Cdk1-Cyclin B activation in controlling cell division.

## RESULTS

### Identification of proteins in the proximity of PKAc using the Turbo-ID

We performed a Turbo-ID to identify novel interactors and substrates of PKA *in ovo*. The Turbo-ID is a proximity-based screening that enables to identify stable and transient interactions between proteins in intact cells^37^. We used catalytic-PKA (PKAc) as bait and fused it with Turbo, a modified and promiscuous biotin ligase that biotinylates proteins within a radius of 10 nm^38^ (Fig. 1A). mRNA coding for either Turbo or Turbo-PKAc were injected in prophase-arrested oocytes and, after 18 hours, oocytes were incubated or not in the presence of biotin. The expression level of Turbo-PKAc is comparable with the endogenous PKAc (Fig. 1B) and Turbo activity is confirmed by an increase in biotinylated proteins in the presence of exogenous biotin (Fig. 1B). Importantly, the expression of Turbo-PKAc increases the phosphorylation of PKAc substrates (Fig. 1B). It is well established that expression of exogenous PKAc inhibits meiotic maturation induced by progesterone^39–42^. Indeed, the expression of Turbo-PKAc inhibits progesterone-induced meiotic maturation, as shown by the absence of oocytes undergoing nuclear envelope breakdown (NEBD) (Sup. Fig. 1A). Additionally, Turbo-PKAc prevents Cdk1 activation in response to progesterone, as shown by the persistence of inhibitory Cdk1 phosphorylation at Y15 and the absence of PP1 phosphorylation at T320, which is a marker of Cdk1 activity (Fig. 1C). The absence of Cdk1 activation also prevents Mos accumulation and MAPK activation (Fig. 1C), which are events occurring downstream Cdk1 activation^29^. Opposingly, Turbo injection does not have any effect on meiosis resumption and on the activation of Cdk1 and the Mos/MAPK pathway (Fig. 1C and Sup. Fig. 1A). Altogether, these experiments show that Turbo-PKAc behaves similarly to PKAc, preventing Cdk1 activation in response to progesterone through the phosphorylation of PKAc physiological substrates.

**Figure 1:**
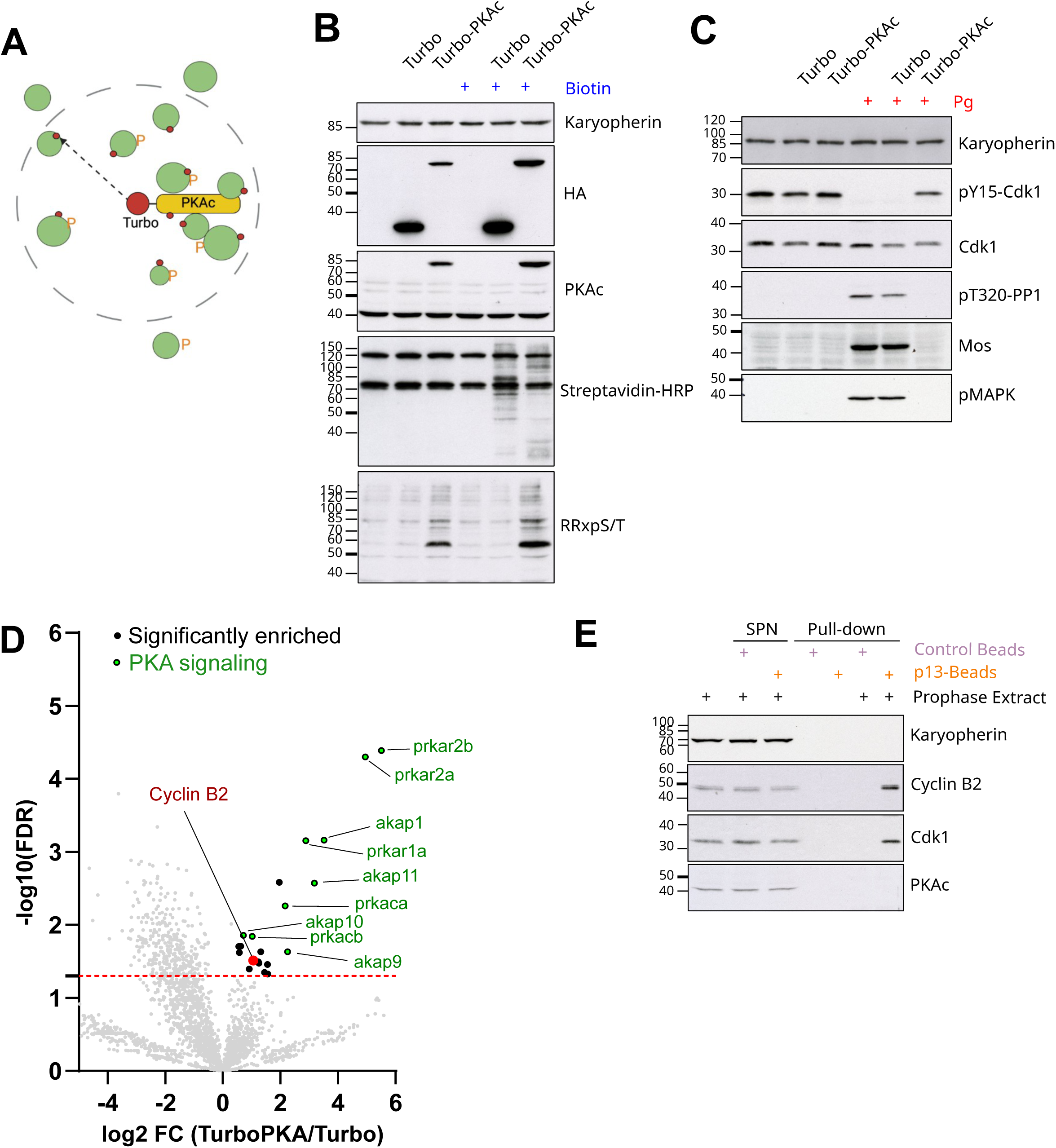
Identification of PKA interactors and substrates by Turbo-ID. (A) Schematic of the Turbo-PKAc proximity-labeling strategy. (B) Prophase-arrested oocytes were injected with mRNA encoding Turbo or Turbo-PKAc. 18 hrs later, they were incubated or not with exogenous biotin for 60 minutes. Lysates were immunoblotted with anti-HA (to detect Turbo/Turbo-PKAc), anti-PKAc, anti-RRxS/T (PKA substrate motif), anti-karyopherin (loading control) and probed with streptavidin–HRP to visualize biotinylated proteins. (C) Prophase-arrested oocytes were injected with mRNA encoding Turbo or Turbo-PKAc. After 18 hrs, oocytes were stimulated with progesterone and collected at NEBD or eNEBD (NEBD equivalent time). Lysates were immunoblotted with anti-Cdk1 pY15, anti-Cdk1, anti-PP1 pT320, anti-Mos, anti-phospho-MAPK (pMAPK), and anti-karyopherin (loading control). (D) Prophase-arrested oocytes were injected with mRNA encoding Turbo or Turbo-PKAc. After 18 hrs, oocytes were incubated with exogenous biotin for 60 min. Biotinylated proteins were recovered by streptavidin pull-down and identified by LC–MS/MS. Four biological replicates were analysed. The volcano plot displays log_2_ fold-change (Turbo-PKAc vs Turbo) on the x-axis versus −log_10_ False Discovery Rate (FDR) on the y-axis. Significantly enriched proteins in Turbo-PKAc are shown in black, known PKA interactors are highlighted in green, and Cyclin B2 is highlighted in red. The red dashed line indicates the FDR threshold of 0.05 (E) Cdk1-Cyclin B complexes were pulled-down with p13-Sepharose beads from prophase-arrested oocyte extracts. Agarose beads were used as control. The oocyte extracts after the pull-down (SPN), and pull-down elution were immunoblotted with anti-Cyclin B2, anti-Cdk1, anti-PKAc, and anti-karyopherin.

The incubation time with exogenous biotin was optimized for the identification of Turbo-PKAc labelled proteins by mass spectrometry (Sup. Fig. 1B-C). High levels of carboxylases were detected independently of the incubation time with biotin in both Turbo-PKAc and Turbo samples (Sup. Fig. 1B-C). Indeed, carboxylases are very abundant enzymes in the oocytes (about 4 μM)^43^ and use biotin as a prostatic factor^44^. As expected, PKA subunits (PKAr and PKAc) and AKAPs (A-kinase Anchoring Proteins 1, 9, 10 and 11) are strongly enriched in Turbo-PKAc compared to Turbo (Fig. 1D and Sup. Fig. 1B-C). Although PKA subunits and AKAPs are already detected after 5 min incubation in exogenous biotin, a 60 min incubation results in their maximal enrichment in Turbo-PKAc over Turbo (Sup. Fig. 1B-C). This data show that Turbo-PKAc expression in oocytes allows to label PKAc known interactors, as PKA subunits and AKAPs, and that their detection is optimal after 60 min incubation in exogenous biotin.

Therefore, the Turbo-ID experiment was performed in four biological replicates using 60 min incubation time with exogenous biotin. The expression of Turbo-PKAc and its activity, as well as the efficiency of the streptavidin pull-down were evaluated (Sup. Fig. 1D) and the biotinylated proteins were identified by mass spectrometry. Notably, more proteins are enriched in Turbo than in Turbo-PKAc, reflecting the higher level of expression of the Turbo construct detected in western blots (Fig. 1B). As expected, PKAc, PKAr and AKAPs proteins are highly enriched in Turbo-PKAc pull-down (Fig. 1D). Surprisingly, we found that Cyclin B2 was significantly enriched in the Turbo-PKAc pull-down (Fig. 1D). Furthermore, Cyclin B2 was consistently enriched across the Turbo-PKAc pull-downs performed during the optimization steps, reinforcing the robustness of its identification in the PKAc proximity-labeling assay (Sup. Fig. 1B).

Detection of Cyclin B2 in proximity of Turbo-PKAc could be due to a stable interaction between the two proteins. To test this hypothesis, a pull-down of Cdk1-Cyclin B complex was performed from prophase oocytes using p13-beads^45,46^. PKAc was not co-precipitated with Cdk1-Cyclin B2, excluding a stable interaction between Cyclin B2 and PKAc in *Xenopus* oocytes (Fig. 1E).

### PKAc phosphorylates Cyclin B2 at S271

Cyclin B2 was identified by Turbo-ID as a PKAc-proximal protein in prophase-arrested oocytes (Fig. 1D). Because no stable interaction between Cyclin B2 and PKAc was detected (Fig. 1E), we considered the possibility that this proximity reflected a transient enzyme– substrate relationship. Notably, *in silico* analysis of the Cyclin B2 sequence reveals the presence of a putative phosphorylation site (R-R-A-S-K) within the Cyclin Box 2 that matches the PKA consensus motif (R/K-R/K-x-pS/T- Φ) (Fig. 2A). To determine whether Cyclin B2 can be phosphorylated by PKAc, Cdk1-Cyclin B2 was purified from prophase oocytes using p13-beads, and the phosphorylation of Cyclin B2 was assessed using an antibody directed against the PKA consensus site (R-R-x-pS/T). Interestingly, endogenous Cyclin B2 is phosphorylated in prophase-arrested oocytes (Fig. 2B). Since endogenous Cyclin B2 phosphorylation is detected by a phospho-PKA substrate motif antibody, we hypothesized that Cyclin B2 is phosphorylated at S271, the only residue matching the PKA consensus motif (Fig. 2A). To assess this, we generated a polyclonal antibody against Cyclin B2 phosphorylated at S271 (Sup. Fig. 2A). 100 ng per oocyte of recombinant Cyclin B2 were added in prophase oocyte extracts, mimicking the endogenous level of the protein (Sup. Fig. 2B). The Δ126-Cyclin B2, corresponding to the human Δ165-Cyclin B, was used because its higher solubility during protein purification ^47^. The phospho-specific anti-pS271 Cyclin B2 antibody detected Cyclin B2 (Fig. 2C), demonstrating that S271 is specifically phosphorylated by a kinase active in prophase-arrested oocytes.

**Figure 2:**
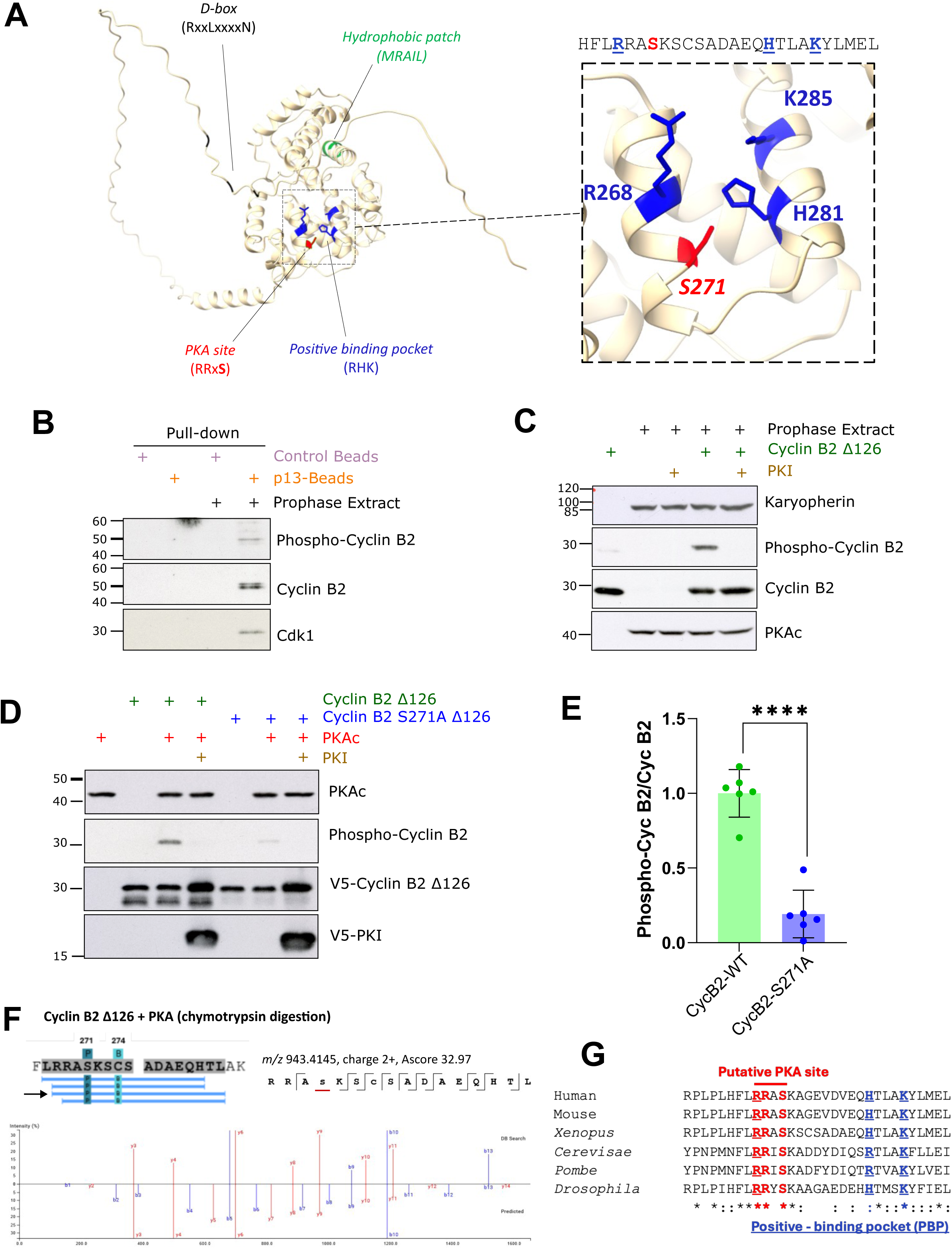
Cyclin B2 is phosphorylated by PKA at S271. (A) Structure of *Xenopus* Cyclin B2 predicted by AlphaFold, highlighting a canonical phospho-PKA consensus motif (R/K– R/K–x–pS/T–Φ) within the Cyclin box 2. S271 in red, the Positive-Binding Pocket in blue, the hydrophobic patch in green, and the D-box in black. The structure was visualized using UCSF ChimeraX. The amino acid sequence of Cyclin B2 surrounding the PBP and PKA phosphorylation consensus motif is reported. (B) Cdk1-Cyclin B2 complexes were isolated from prophase-arrested oocyte extracts using p13-beads. Agarose beads were used as control. Pull-down elutions were immunoblotted with anti-RRxpS/T (phospho-Cyclin B2), anti-Cyclin B2 and anti-Cdk1 antibodies. (C) Prophase extracts were supplemented or not with the PKA inhibitor PKI. After a 15 min incubation, 100 ng of recombinant Δ126-Cyclin B2 was added for 60 min. Lysates were immunoblotted with anti-karyopherin (loading control), anti-pS271 Cyclin B2 (phospho-Cyclin B2), anti-Cyclin B2 and anti-PKAc antibodies. (D) Commercial PKAc was incubated or not with V5-PKI for 30 min. V5-Δ126-Cyclin B2 WT or V5-Δ126-Cyclin B2-S271A were added for 30 min. Reactions were immunoblotted with anti-PKAc, anti-RRxpS/T (phospho-Cyclin B2), and anti-V5 (to detect V5-Δ126-Cyclin B2 and V5-PKI) antibodies. (E) Quantification of Cyclin B2 phosphorylation (ratios between RRxpS/T signals and V5-Cyclin B2) from six biological replicates of the experiment described in panel D. Mean and S.E.M. are displayed. A t-test was performed to assess the statistical significance. ****: p-value < 0.0001. (F) V5-Δ126-Cyclin B2 WT was incubated with commercial PKAc for 30 minutes. Phosphorylation sites were analysed by MS. MS/MS fragmentation spectrum of the RRASKSCSADAEQHTL peptide carrying a phosphorylation at S4, corresponding to S271 in the Cyclin B2 sequence. Site localization confidence was assessed using an Ascore, defined as an ambiguity score of −10×log⁡10(p). The p-value reflects the probability that the peptide match occurred by chance (Ascore = 20 for p = 0.01). (G) Alignment of Cyclin B2 sequences from different species. The PKA consensus and the Positive-Binding Pocket (PBP) are indicated in red and blue respectively.

To investigate whether PKA is the kinase responsible on the phosphorylation of Cyclin B2 at S271, prophase oocytes extracts were supplemented with PKI, a strong inhibitor of PKA^6,29^. Cyclin B2 phosphorylation at S271 was prevented by the preincubation of the extract with PKI (Fig. 2C). Arpp19-GST phosphorylation at S109 by PKAc was used as a control and displays a similar behaviour to that of Cyclin B2^28,29^(Sup. Fig. 2C). This result demonstrates that Cyclin B2 is phosphorylated at S271 by PKAc in prophase oocyte extracts.

We next assessed whether PKAc directly phosphorylates Cyclin B2 *in vitro*. Incubation of recombinant Cyclin B2 with PKAc induced a strong phosphorylation signal, which was abolished by PKI and markedly reduced by mutation of S271 to alanine (S271A) (Fig. 2E–F). Mass spectrometry further confirmed phosphorylation of Cyclin B2 at S271 under these conditions (Fig. 2F). Together, these results demonstrate that PKAc directly phosphorylates Cyclin B2 at S271.

Notably, the site of PKA phosphorylation is well conserved in Cyclin B1, but not in Cyclin B3 and any A-type Cyclins, in both *Xenopus* and human (Sup. Fig. 2D). Hence, PKAc ability to phosphorylate human Cyclin B1 and Cyclin A2 was investigated *in vitro*. A Cdk1 inhibitor, Cip1, was used to exclude any contribution of Cdk1 activity to the assay. PKAc phosphorylates human Cyclin B1, but not human Cyclin A2, and this phosphorylation is suppressed by PKI (Sup. Fig. 2E). Importantly, PKA consensus site of phosphorylation (R-R-x-S-K) of Cyclin B2 is highly conserved across eukaryotes from yeast to human (Fig. 2G), suggesting a conserved role of this phosphorylation.

### Phosphorylation of Cyclin B2 at S271 suppresses its ability to induce meiosis resumption

It was shown that Cyclin B and Cyclin A injection in prophase oocytes induces meiotic maturation independently of progesterone stimulation^48,49^. Since PKA maintains the prophase arrest through the phosphorylation of its substrates, we evaluated whether PKA phosphorylation of Cyclin B2 affects its ability to induce meiotic resumption in *Xenopus* oocyte. Hence, we produced phospho-mimetic (S271D) and phospho-null (S271A) mutants of Cyclin B2 (Fig. 3A) and mRNA encoding these constructs were injected in prophase-arrested oocytes. Cyclin A was used as a positive control. 18 hrs after microinjection, the oocytes were scored for the presence of the white spot marking NEBD and Cdk1 activation, and collected for western blot analysis (Fig. 3B-C). As expected, the injection of wild-type Cyclin B2 (Cyclin B2-WT) induced meiotic maturation in the absence of progesterone (Fig. 3B-C). Indeed, Cdk1 was activated in oocytes undergoing NEBD as marked by the dephosphorylation of Cdk1 at Y15, the phosphorylation of its substrate, PP1, at T320, and phosphorylation of MAPK (Fig. 3B). A similar phenotype was observed with Cyclin B2-S271A. Interestingly, Cyclin B2-S271D completely lost the ability to induce NEBD and Cdk1 activation, even though it was expressed at the same levels of Cyclin B2-WT or Cyclin B2-S271A (Fig. 3B-C). These findings demonstrate that the phosphorylation state of S271 controls the ability of Cyclin B2 to induce meiotic maturation.

**Figure 3:**
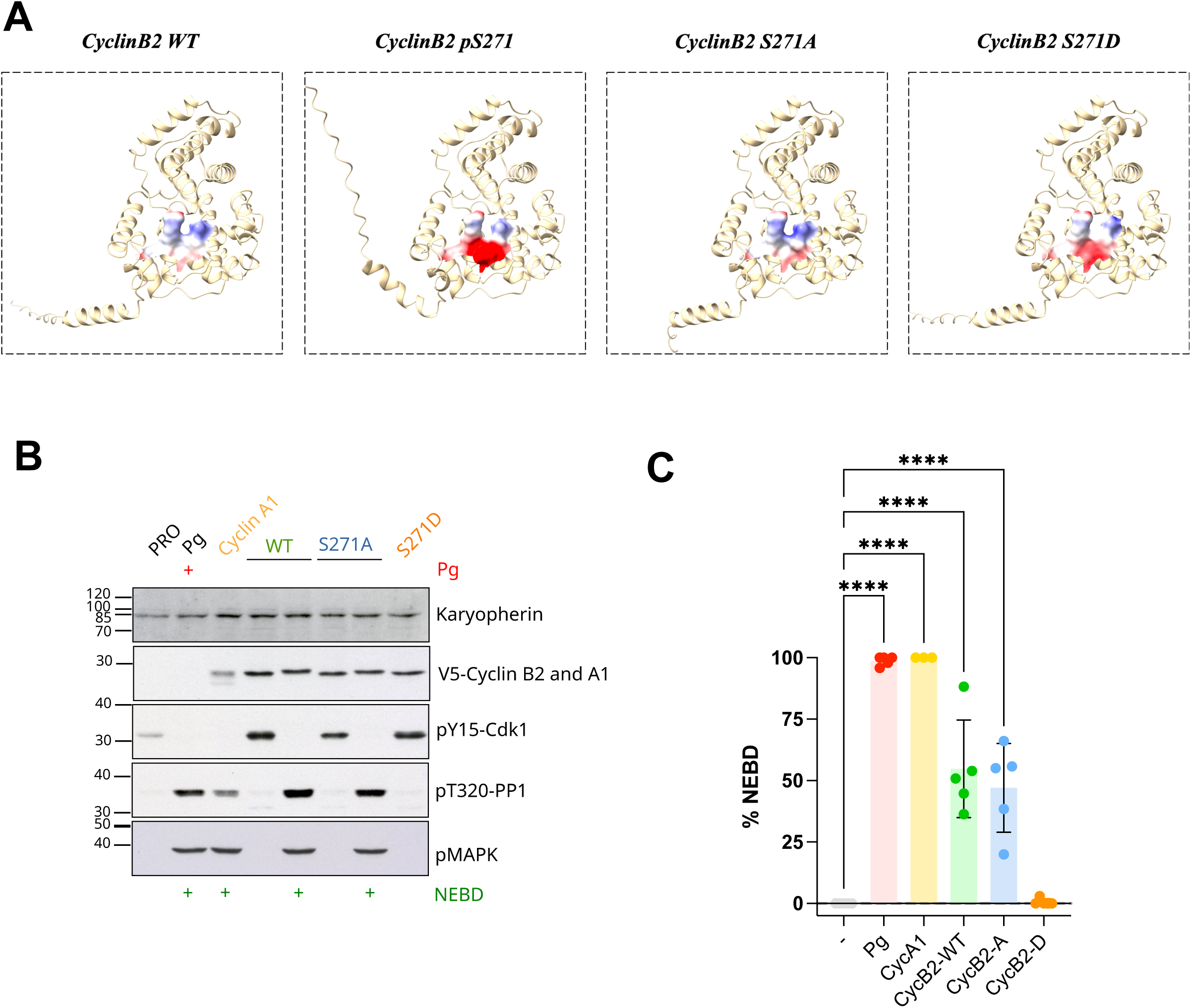
The phosphorylation of Cyclin B2 at S271 impairs its ability to promote meiotic maturation. (A) AlphaFold model of Δ90-Cyclin B2 (WT, phosphorylated at S271 -pS271-, S271A, or S271D). The electrostatic surface potential of the residues composing the PBP (R268, H281 and K285) and S271 is shown. Positive and negative electrostatic potential are represented in blue and red, respectively. AlphaFold predicted structures were visualized and the electrostatic surface potentials were calculated using UCSF ChimeraX. (B-C) Prophase-arrested *Xenopus* oocytes were injected with mRNA encoding V5-tagged Δ90-Cyclin B2 (WT, S271A, or S271D) or V5-Cyclin A1. After 18 hrs, oocytes were scored for NEBD (white spot) (C) and collected for western blot (B). (B) Western blots with anti-pY15-Cdk1, anti-pT320-PP1, anti-phospho-MAPK (pMAPK), anti-V5 (to detect exogenous Cyclins) and anti-karyopherin (loading control) antibodies. (C) Percentage of NEBD 18 hrs after mRNA microinjection from 5 biological replicates; data are shown as mean ± S.E.M. An ANOVA was performed to assess the statistical significance. ****: p-value < 0.0001.

### PKAc phosphorylation of Cyclin B2 does not affect its stability or its ability to bind Cdk1

It was previously shown that the Cdk activator RINGO/Speedy undergoes ubiquitin– proteasome-mediated degradation controlled by PKA during the prophase-arrest of *Xenopus* oocytes^34^. Additionally, we have previously shown that Cyclin B accumulates in response to PKA downregulation without any change in its translation rate^29^, suggesting that it is regulated at the level of its turnover. Therefore, we asked whether PKAc-dependent phosphorylation of Cyclin B2 limits its accumulation by promoting its degradation, which could explain why the phospho-mimetic mutant fails to trigger meiotic resumption. mRNAs encoding full length wild-type Cyclin B2, phospho-mimetic or phospho-null mutants were injected in prophase oocytes, and their accumulation was assessed after 24h (Fig. 4A). mRNA encoding MBP-GST was co-injected with the Cyclin B2 mRNA, as a loading control. No difference in protein accumulation was observed, suggesting that the phosphorylation does not modulate Cyclin B2 turnover (Fig. 4A-B).

**Figure 4:**
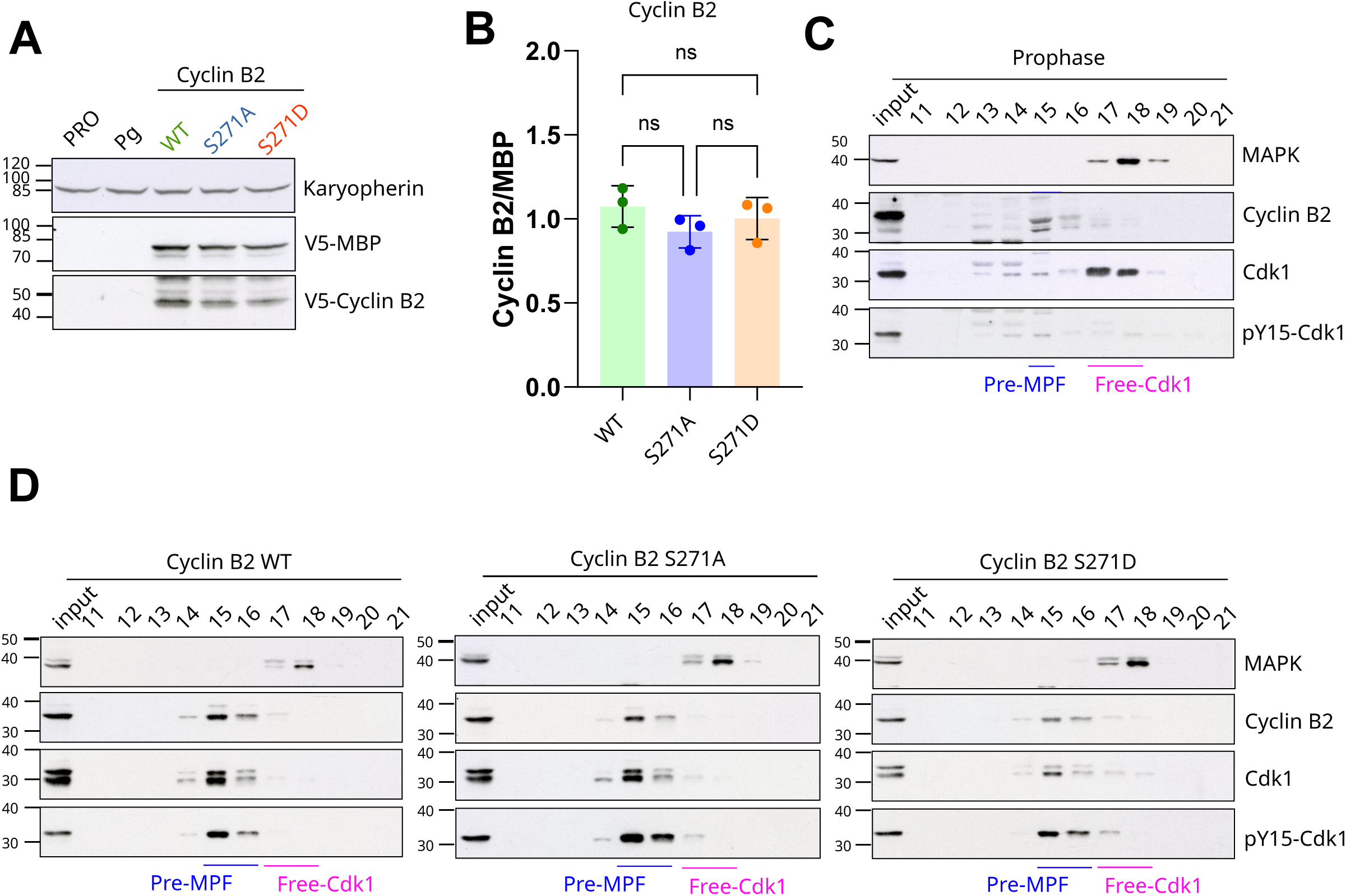
Phosphorylation of Cyclin B2 at S271 does not affect its stability or its interaction with Cdk1. (A) Prophase-arrested *Xenopus* oocytes were injected with mRNA encoding full length V5–Cyclin B2 (WT, S271A, or S271D) together with V5-MBP-GST. Oocytes were collected 24 hrs later, lysed and immunoblotted with anti-V5 (to detect exogenous Cyclin B2 and MBP-GST) and anti-karyopherin (loading control) antibodies. (B) Quantification of Cyclin B2 accumulation from three independent biological replicates of the experiment shown in (A) (V5–Cyclin B2 normalized to V5–MBP-GST); data plotted as mean ± S.E.M.. An ANOVA was performed to assess the statistical significance. (C) Prophase oocyte extracts were fractionated by Superose 12 size exclusion chromatography. Total extracts (input) and fractions 11 to 21 were immunoblotted with antibodies against Cyclin B2, Cdk1, pY15-Cdk1 and MAPK (Erk) as control. (D) Extracts from prophase oocytes expressing V5- Δ90-Cyclin B2, either wild type (WT), or mutated (S271A or S271D) were fractionated through a Superose 12 size exclusion chromatography. Total extracts and fractions 11 to 21 were immunoblotted as in (C).

Alternatively, S271 phosphorylation may impede Cyclin B2 binding to its partner Cdk1. The ability of wild-type Cyclin B2 and phospho-mutants to bind Cdk1 *in ovo* was assessed by size exclusion chromatography. As previously reported^50^, Cdk1 is present in prophase-arrested oocytes in two distinct pools: a minor one associated with Cyclin B2 (fraction 15), and a predominant monomeric pool (fractions 17-18)(Fig. 4C). Injection of mRNAs encoding wild-type Cyclin B2 in prophase-arrested oocytes led to an increase in the amount of Cdk1-Cyclin B2 complexes (fractions 15-16) (Fig. 4D). This expected result is due to the binding of exogenous Cyclin B2 to the pool of monomeric Cdk1, leading to its reduction in fractions 17-18 (Fig. 4D). The formation of these new Cdk1-Cyclin B2 complexes is also confirmed by an increase in the level of Cdk1 phosphorylation at Y15 that occurs only after Cyclin B binding^51^, due to the high activity of Myt1 in prophase-arrested oocytes (Fig. 4D)^52^. The elution profile of MAPK (fraction 18) was used as an internal control to confirm the reproducibility of the fractionation procedure (Fig. 4C-D). Interestingly, oocytes injected with mRNAs encoding either Cyclin B2-S271A or Cyclin B2-S271D mutant displayed an elution profile indistinguishable from that of wild-type Cyclin B2 (Fig. 4D). These results demonstrate that Cyclin B2 can efficiently bind Cdk1 *in vivo* regardless of the phosphorylation status of S271.

### Phosphorylated Cyclin B impairs Cdk1 multi-phosphorylation of its substrates

Having excluded any effect of S271 phosphorylation of Cyclin B2 on its stability or its Cdk1 binding, we next asked whether this phosphorylation event could prevent Cdk1 activation. Cdk1-Cyclin B complexes phosphorylate their substrates through two complementary mechanisms: recognition of a canonical Cdk1 consensus motifs by the kinase subunit, Cdk1, and substrate docking mediated by Cyclin B itself (Fig. 5A). We therefore tested whether PKA phosphorylation of Cyclin B selectively affects one of these modes of substrate phosphorylation. Since the phosphorylation site is conserved between Cyclin B2 and Cyclin B1 (Sup. Fig. 2D), and PKAc phosphorylates Cdk1-Cyclin B1 (Sup. Fig. 2E), we used a commercially available active Cdk1-Cyclin B1 complex. Cdk1-Cyclin B1 was pre-phosphorylated or not by PKAc. The effect of the phosphorylation on Cdk1 activation was evaluated by its ability to phosphorylate PP1, a well-established Cdk1 substrate that is directly recognized by Cdk1^53,54^. As expected, Cdk1-Cyclin B1 phosphorylates PP1 at T320 (Fig. 5B). Interestingly, pre-phosphorylation of Cyclin B1 by PKAc does not alter Cdk1 ability to phosphorylate PP1 (Fig. 5B-C).

**Figure 5:**
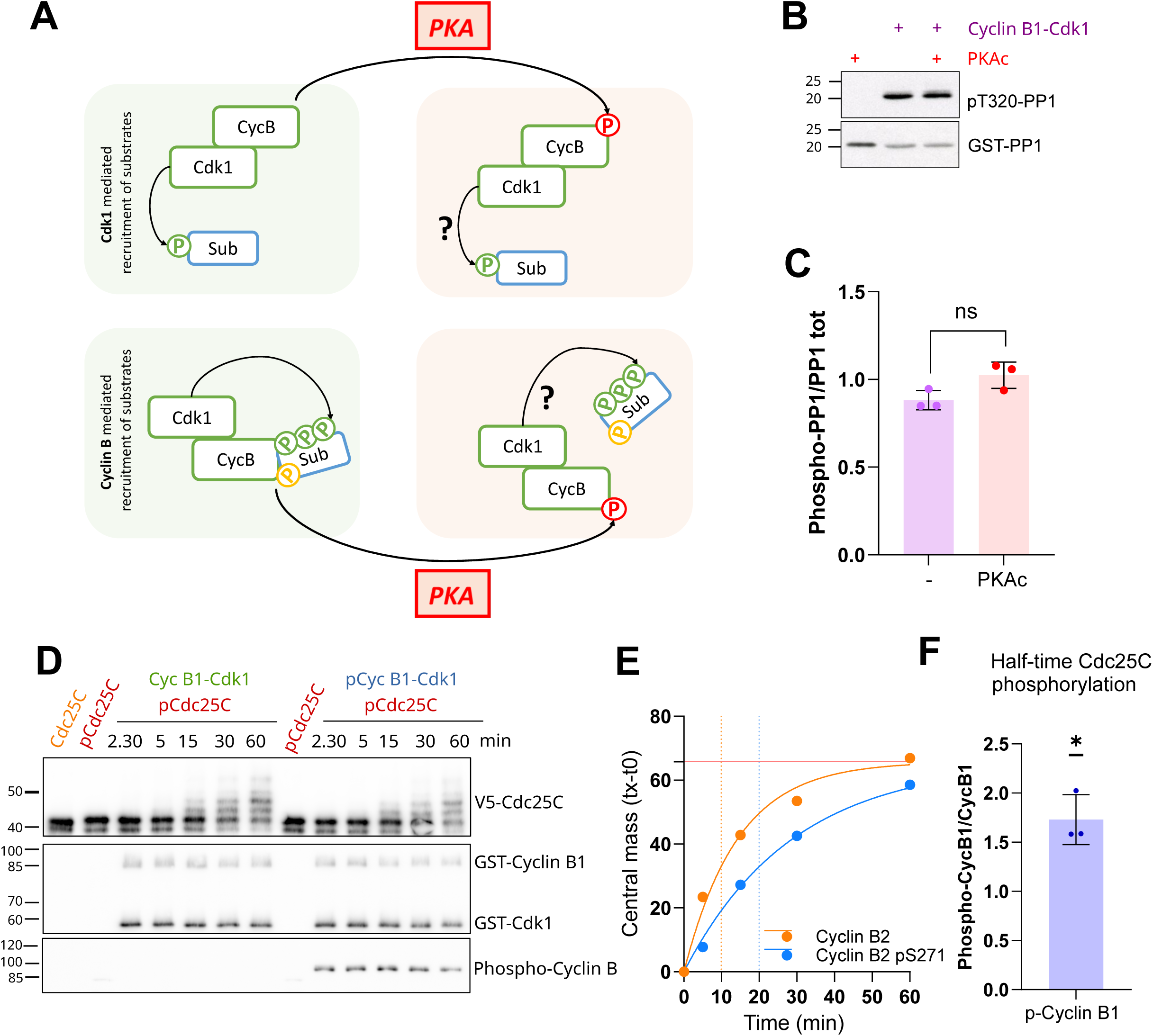
PKA phosphorylation of Cyclin B at S271 impairs its ability to mediate multi-site phosphorylation. (A) Schematic representation of the substrate recognition mediated by either Cdk1 or Cyclin B. (B) Human GST-Cyclin B1-Cdk1 were incubated or not in the presence of PKAc for 30 min. The Cdk1 substrate, PP1-GST, was then added for 30 min. Reactions were immunoblotted with anti-pT320-PP1 and anti-GST-HRP antibodies. (C) Quantification of PP1 phosphorylation at T320 (ratio between anti-pT320-PP1 and GST-HRP) from three biological replicates of the experiment shown in (B). Mean and S.E.M. are displayed. A t-test was performed to assess the statistical significance. (D) Recombinant V5-Cdc25C (sequence 89-342, pCdc25C) was incubated first in the presence of commercial PKAc. V5-PKI was added for 30 min to stop the reaction. Commercial Cyclin B1-Cdk1 was incubated (pCycB/Cdk1) or not (CycB/Cdk1) for 30 min in the presence of PKAc. pCdc25C was then added. Samples were collected at 2.30, 5, 15, 30, or 60 min. Reactions were immunoblotted with antibodies against V5 (to detect Cdc25C), GST-HRP (to detect Cyclin B1 and Cdk1) and RRxpS/T (to detect Cyclin B1 phosphorylation). (E) The centre of mass for each upshift in migration of V5-Cdc25C was quantified as described in the methods. (F) The half-time of three independent replicates was expressed as ratio of non-phosphorylated Cyclin B1 half-time. Single sample t-test with null hypothesis of 1 was performed. *: p-value < 0.05.

We then investigated the effect of Cyclin B phosphorylation on the phosphorylation of substrates that are recruited through Cyclin B. Interestingly, the PKA phosphorylation site of Cyclin B2 resides inside the recently identified PBP^24,25^, which docks previously phosphorylated substrates and primes their subsequent phosphorylations at non-consensus Cdk1 sites (Fig. 2). Therefore, we hypothesized that PKA phosphorylation at S271 of Cyclin B2 could alter this mechanism by supressing the polar charge of this domain (Fig. 3A). We examined this possibility using Cdc25C, because it undergoes multi-site phosphorylation by Cdk1-Cyclin B and its interaction with Cdk1-Cyclin B has been proposed to involve residues within the Cyclin B PBP, including the RRASK motif^55^. Because Cdc25C is also a PKA substrate in prophase-arrested oocytes^26^, recombinant Cdc25C was first pre-phosphorylated by PKAc. Cdk1-Cyclin B1 complexes, pre-phosphorylated or not by PKAc, were then assayed for their ability to phosphorylate Cdc25C over time. Cdk1-Cyclin B progressively phosphorylated Cdc25C, as visualized by a gradual decrease in its electrophoretic mobility on SDS-PAGE (Fig. 5D). To quantify this shift, the displacement of the Cdc25C signal was measured throughout the kinase assay by calculating the center of mass of the band intensity profile, allowing to determine the half-time of Cdc25C phosphorylation (Fig. 5E). This analysis showed that phosphorylation of Cyclin B1 by PKAc slows Cdc25C phosphorylation (Fig. 5D–F). These results show that PKA phosphorylation of Cyclin B limits the ability of Cdk1–Cyclin B to promote multisite phosphorylation of Cdc25C, thereby dampening a key step in the Cdk1 activation network and explaining why the phospho-mimetic mutant fails to induce meiosis resumption (Fig. 3B-C).

## DISCUSSION

For more than five decades, it has been established that PKA activity maintains prophase arrest in vertebrate oocytes and that PKA inhibition is required for meiotic resumption. Despite this long-standing knowledge, the few PKA substrates identified to date have not provided a mechanistic explanation for how their dephosphorylation could lead to Cdk1 activation. Our findings address this longstanding question. Here, we identify Cyclin B as a direct target of PKA in vertebrate oocytes. Using a PKAc-based proximity-labeling approach in prophase-arrested *Xenopus* oocytes, we found that Cyclin B2 is enriched in the proximity to PKAc. Although Cyclin B2 does not stably associate with PKAc, it contains a conserved PKA consensus site within the Cyclin Box, centered on S271. We demonstrate that PKAc directly phosphorylates Cyclin B2 at this site both *in vitro* and in oocyte extracts. Functionally, phospho-mimetic substitution of S271 abolishes the ability of Cyclin B2 to induce meiotic maturation and Cdk1 activation, without affecting Cyclin B2 accumulation or its association with Cdk1. Instead, phosphorylation of Cyclin B selectively impairs multisite phosphorylation of Cdc25C by Cdk1-Cyclin B. Together, these findings identify Cyclin B as a direct PKA-regulated component of the Cdk1 activation network and uncover an essential mechanism by which PKA controls entry into meiotic division.

These results provide a mechanistic explanation for the long-standing observation that Cyclin B-induced meiotic maturation remains sensitive to PKA activity. In *Xenopus* oocytes, downregulation of PKA is required for the protein-accumulation program that generates the initial amount of active Cdk1, but PKA also restrains the subsequent auto-amplification loop that converts this initial activity into full Cdk1 activation. Our data suggest that phosphorylation of Cyclin B contributes to this second lock, together with the PKA-dependent regulation of Cdc25C. In this model, PKA does not simply prevent the accumulation of Cyclin B or its binding to Cdk1; rather, it limits the ability of pre-existing Cdk1-Cyclin B complexes to phosphorylate key regulators of their own activation. Thus, PKA maintains prophase arrest through a multilayered mechanism, acting both upstream of Cdk1 activation and directly on the Cdk1-Cyclin B complex that drives M-phase entry.

An important point is that both Cyclin B1 and Cyclin B2 contain the conserved PKA site. In prophase-arrested *Xenopus* oocytes, Cyclin B2 is more abundant than Cyclin B1, which may explain why Cyclin B2 was identified in the PKAc proximity-labeling screen. However, our *in vitro* assays show that Cyclin B1 is also phosphorylated by PKA, indicating that this regulatory mechanism is not restricted to Cyclin B2. This raises the possibility that phosphorylation of pre-existing Cyclin B2 mainly acts as a safeguard to prevent premature activation of the pre-MPF pool during prophase arrest, before the hormonal signal has been received. In this context, Cyclin B phosphorylation may cooperate with the inhibitory Cdk1 phosphorylation by Myt1 to maintain Cdk1-Cyclin B complexes in an inactive state. Consistent with this idea, pharmacological inhibition of Myt1 with PD0166285 only inefficiently induces meiotic maturation, suggesting that additional mechanisms restrict the activation of MPF^52^.

Upon PKA downregulation, newly synthesized Cyclin B1 accumulates and, unlike pre-existing Cyclin B2, is not phosphorylated at S271. Hence, Cyclin B1 may contribute to Cdk1 activation without requiring dephosphorylation of the pre-existing Cyclin B2 pool. Therefore, Cyclin B2 phosphorylation by PKA may function as a guardian mechanism that restrains activation of stored Cdk1-Cyclin B complexes. In this model, Cyclin B1 molecules accumulating following PKA inhibition would escape this phosphorylation, providing a threshold activity independently of Cyclin B2 dephosphorylation, to trigger the auto-amplification loop.

Our data identify Cdc25C as a target whose multisite phosphorylation by Cdk1-Cyclin B is sensitive to Cyclin B phosphorylation by PKA. This is particularly relevant because Cdc25C is both a PKA-regulated protein in prophase and a central component of the Cdk1 auto-amplification loop. PKA phosphorylation of Cyclin B could therefore restrain Cdk1 activation indirectly by reducing the efficiency of Cdk1-Cyclin B to phosphorylate and activate Cdc25C. Whether this regulation is specific to Cdc25C or reflects a broader control of Cdk1 substrate selection remains an important open question. Multisite phosphorylation is a common feature of many Cdk1 substrates, and Cyclin B docking surfaces are increasingly recognized as major determinants of substrate specificity, as recently shown for the APC components^25^. Future work should define how the different substrate-recruitment modules of Cyclin B, including the hydrophobic patch and the positively charged phosphate-binding pocket, cooperate to recruit distinct classes of substrates. It will also be important to understand how these Cyclin B-dependent docking mechanisms are integrated with Cks-dependent recognition of phosphorylated substrates during the progressive phosphorylation of M-phase regulators.

Interestingly, the PKA phosphorylation site identified in Cyclin B is absent from Cyclin A proteins. This further supports the notion that, despite both being Cdk1-associated cyclins, Cyclin A and Cyclin B have distinct roles in promoting M-phase entry. Notably, Cyclin A, which accumulates only later during meiotic resumption, does not contribute to the initiation of meiotic division. Conversely, in somatic cells, Cyclin A has been proposed to promote mitotic entry by escaping the major inhibitory control imposed on Cyclin B–Cdk1 complexes by the Myt1/Wee1 kinases. Our findings reveal that Cyclin A also lacks the PKA-regulated inhibitory mechanism identified in Cyclin B, further highlighting how distinct Cyclin classes have evolved different regulatory properties to control the G2-M transition.

Finally, the strong conservation of the PKA phosphorylation site across eukaryotic Cyclin B proteins suggests that this regulatory mechanism may extend beyond *Xenopus* oocyte meiosis. We show that human Cyclin B1 is phosphorylated by PKA *in vitro*, supporting the idea that this regulatory site has been functionally conserved across vertebrates. Notably, the motif is also present in Cyclin B proteins from non-vertebrate species in which meiotic arrest is not controlled by PKA. This suggests that phosphorylation of Cyclin B may have a broader function in modulating Cdk1-Cyclin B activity beyond meiosis. Interestingly, Cdc25C phosphorylation by PKA occurs on the same site as that phosphorylated by Chk1 during the DNA damage checkpoint^56^. S271 also lies within a putative Chk1 consensus motif (K/RxxS/T). Whether S271 phosphorylation by kinases other than PKA, such as Chk1/2, contributes to the G2/M arrest following DNA damage in somatic cells remains an intriguing question for future studies.

## METHODS

### Animals

Adult female *Xenopus laevis* were sourced from the Centre de Ressources Biologiques Xénopes (CNRS, France) and housed in the aquatic animal facility of the IBPS, following French regulatory standards (Animal Facility Agreement: #A75-05-25). All animal experiments were conducted under protocols approved by the French Ministry of Higher Education and Research (authorization APAFIS #45718-2025021111055055v2).

### Xenopus oocytes

Oocytes were harvested from adult, non-hormonally primed *Xenopus laevis* females. Animals were anesthetized by immersion for 30 min in 1 g/L MS-222 (Sigma, E10521) buffered with sodium bicarbonate. Ovarian lobes were removed and enzymatically dissociated in M buffer (10 mM HEPES, pH 7.8; 88 mM NaCl; 1 mM KCl; 0.33 mM Ca(NO₃)₂; 0.41 mM CaCl₂; 0.82 mM MgSO₄) containing 0.4 mg/mL dispase II (Roche, 04942078001) for 3 hrs, followed by 0.4 mg/mL collagenase (Sigma, C9891) for 1 hr. After digestion, oocytes were extensively washed with 2 L of M buffer to remove residual enzymes. The largest oocytes, corresponding to stage VI fully-grown oocytes^57^. Maturation was induced with 1 µM progesterone. Oocytes exhibiting a white spot at the animal pole resulting from the first pigment rearrangement were scored as having undergone NEBD.

### Western blot

Oocytes were lysed in extraction buffer (EB) (80 mM β-glycerophosphate, 20 mM EGTA, 15 mM MgCl_2_, pH 7.3) with a ratio of 1 oocyte in 10 µl, loaded in Laemmli gels for electrophoresis and transfer onto nitrocellulose membranes (Amersham, 10600015) with semi-dry apparatus of transfer^58^. After overnight incubation with primary antibodies at 4°C, membranes were incubated with appropriate horseradish peroxidase-labelled secondary antibodies (Jackson Immunoresearch) for 2 hrs at room temperature. The signal was revealed with chemiluminescence (BioRad, Clarity^TM^ Western ECL Substrate, 1705061).

The following primary antibodies diluted in PBS, 3% BSA, 0.05% NaN_3_ were directed against: HA (1:5000, mouse, Sigma H9658), PKAc (1:10.000 for the detection of the endogenous protein and 1:20.000 for the kinase assays, rabbit, Abcam Ab26322), Karyopherin (1:3000, goat, Santa Cruz sc-1863), Streptavidin-HRP (1:20000, Pierce^TM^ High Sensitivity HRP-conjugated, Thermo Scientific 21130), RRxpS/T (1:5000 or 1:10.000 for *in vitro* kinase assays, rabbit, Cell Signaling 9624), pY15-Cdk1 (1:1000, rabbit, Cell Signaling 9111), Cdk1 (1:3000, mouse, Invitrogen MA-91598), pT320-PP1 (1:5000 for the detection of the endogenous protein, 1:20.000 for the kinase assay, rabbit, Abcam ab62334), Mos (1:500, rabbit, Santa Cruz Biotechnology Sc-86), pMAPK (1:2000, mouse, Cell Signaling 9106), Cyclin B2 (1:1000, mouse, Abcam ab18250 or 1:50, goat^50^), phospho-S271-Cyclin B2 (1:50.000, rabbit), V5 (1:10.000 for mRNA injection in oocytes or 1:20.000 for *in vitro* kinase assays, mouse, Invitrogen 46–0705), GST-HRP (1:50.000 overnight, 1:10.000 for PP1-KA, HRP-conjugated, RPN 1236V), Erk1/2 -MAPK- (1:2000 each, rabbit, Santa Cruz Biotechnology C-16 and C-14), pS109-Arpp19 (1:400.000 for the kinase assays, rabbit^28^).

### Turbo-ID

Oocytes were micro-injected with either mRNA coding for Turbo-PKAc (30 ng) or Turbo (15 ng). After 18 hrs, oocytes were incubated with 50 µM of exogenous biotin (B4639 Sigma-Aldrich) for 60 min. 50 oocytes per condition were lysed in EB and supplemented with buffer A (2M Tris pH 7.4, 1M NaCl, 20% NP40, 2% SDS). Extracts were pre-cleaned with pre-washed Dynabeads protein A (10002D Invitrogen) for 2 hrs at 4°C. The pull-down was performed with 120 μl of pre-washed Dynabeads^TM^ MyOne^TM^ Streptavidin C1 beads (65001 Invitrogen) at 4°C for 18 hrs under rotation. Beads were washed under rotation with the following buffers: twice with buffer A for 5 min at 25°C, three times with buffer B (50 mM Tris pH 7.4 and 8 M urea) for 5 min at 25°C, once with buffer A for 5 min at 25°C. Beads were stored in MilliQ water at -80°C until further analysis.

### LC–MS/MS analysis and data processing

Beads from pull-down experiments were digested overnight at 37 °C in 20 µL of 50 mM NH₄HCO₃ containing 1 µg sequencing-grade trypsin/Lys-C mix. Peptides were loaded onto Evotips (Evosep, Odense, Denmark) and desalted according to the manufacturer’s instructions prior to LC–MS/MS analysis. Peptides were analysed on a timsTOF Pro 2 mass spectrometer (Bruker Daltonics, Bremen, Germany) coupled to an Evosep One system. Turbo-PKAc samples were acquired using the 30 samples-per-day (30SPD) method with a 44-min gradient and 48-min total cycle time on a C18 analytical column (0.15 × 150 mm, 1.9 µm beads, EV-1106) maintained at 40 °C and operated at 500 nL min⁻¹. Solvent A was H₂O/0.1% formic acid (FA) and solvent B was acetonitrile/0.1% FA. The instrument was operated in PASEF mode with a 1.3-s cycle time, and MS/MS spectra were acquired over an m/z range of 100–1700.

Samples from the *in vitro* PKAc phosphorylation assay were digested with chymotrypsin and analysed using the 40 samples-per-day (40SPD) Whisper Zoom method with a 32-min gradient and 36-min total cycle time. Peptides were separated on a C18 column (0.075 × 150 mm, 1.7 µm beads, Aurora Elite CSI, IonOpticks) maintained at 50 °C and operated at 200 nL min⁻¹. Raw MS data were processed using PEAKS Online 11 (build 1.9; Bioinformatics Solutions Inc.) and searched against the *Xenopus* laevis protein database (downloaded January 2023; 25,839 entries). To reduce redundancy, L and S alleles were merged together under a single accession number. Parent and fragment mass tolerances were 20 ppm and 0.05 Da, respectively. Specific digestion was selected with up to two missed cleavages permitted. Oxidation (M), deamidation (NQ), phosphorylation (STY) and acetylation (protein N-term) were set as variable modifications, and half cysteine (C) as a fixed modification. Peptide and protein group identifications were both filtered at a 1% false discovery rate.

Label-free quantification for Turbo-PKAc samples was performed using the PEAKS quantification module with a mass tolerance of 10 ppm, a CCS tolerance of 0.02 and a retention time shift tolerance of 0.5 min for match-between-runs. Protein abundance was calculated using the Top All Peptides method and total ion current (TIC) normalization was applied. Multivariate analysis of protein abundances was performed using Qlucore Omics Explorer 3.9 (Qlucore AB, Lund, Sweden). A positive threshold value of 1 was specified to enable a log2 transformation of abundance data for normalization *i.e.* all abundance data values below the threshold will be replaced by 1 before transformation. Differential proteins between groups were identified using a two-sided Student’s t-test.

### Cloning and vectors

Cloning was performed using assembly (NEBuilder^®^ HiFi DNA Assembly Master Mix) or overlap extension PCR (NEB, Q5^®^ High-Fidelity 2X Master Mix) and analysed by sanger sequencing (GenewizAzenta). The following plasmids were used: GST-XenArpp19-WT; GST-XenCip1 (NM_001094464.1); 6xHis-v5-PKIg; MBP-GST-V5 as in ^29^; 6xHis-HA-Turbo-PKAc; 6xHis-V5-Δ126-XeCCNB2; 6xHis-V5-Δ126-XeCCNB2-S271A; T3-6xHis-V5-XeCCNB2.L; T3-6xHis-V5-XeCCNB2.L_S271A; T3-6xHis-V5-XeCCNB2.L_S271D; T3-6xHis-V5-CCNA1.L; ORF-pGEX-6P-2original_6xHis_Cdc25C.S-fragment.

### mRNA and protein production

Template for *in vitro* transcription were produced with primers and vector reported in Table 1. mRNA was *in vitro* produced using the mMESSAGE mMACHINE T3 Transcription Kit (Ambion, AM1348) and polyadenylated (150-200 nt) with Poly(A) Tailing Kit (Ambion, AM1350). Messengers were purified using MEGAclear Kit (Ambion, AM1908) and correct length and polyadenylation were assessed by electrophoresis. mRNA concentration was evaluated by NanoDrop. The following quantities of mRNA were injected in each oocyte: Turbo-PKAc 30 ng, Turbo 15 ng, Δ90-WT-CyclinB2, Δ90-S271A-Cyclin B2, Δ90-S271D-Cyclin B2 and Δ90-Cyclin A1 for testing the effect of the phosphorylation of Cyclin B2 ability to induce maturation: 25 ng; WT-Cyclin B2, S271A-Cyclin B2 and S271D-Cyclin B2: 25 ng; MBP-GST-V5 for assessing the stability of Cyclin B2: 2.5 ng; Δ90-Cyclin B2, Δ90-S271A-Cyclin B2 and Δ90-S271D-Cyclin B2 for testing Cyclin B2 binding to Cdk1: 25 ng.

Recombinant PKI and Cip1 were expressed and purified as described in^29^ and GST-Arpp19 as in^28^. Δ126-Cyclin B2 and Δ126-S271A-Cyclin B2 were obtained as similarly described in ^47^. Recombinant Cdc25C fragment (89-342) was produced in *Escherichia coli* by autoinduction (Studier 2005) and purified using TALON-Cobalt beads column (635507, Takara Bio) for 6XHIS-tagged protein as previously described in^28,54^.

The recombinant proteins were produced and purified on glutathione-agarose column (G4510, Sigma) for GST-tagged protein or on TALON-Cobalt beads column (635507, Takara Bio) for 6XHIS-tagged protein as previously described.

### Pull-down

100 prophase oocytes were lysed in buffer C (80 mM β-glycerophosphate, 10 mM EDTA, 30 mM NaCl, anti-proteases - cOmplete^TM^ Sigma Aldrich, 10 µM okadaic acid OA - Enzo Life Sciences or Sigma, 42 µg/ml NaF and 210 µg/ml Na_3_VO_4_, pH 7.4). The extract was supplemented with buffer D (375 mM NaCl, 0.15% Triton, anti-proteases - cOmplete^TM^ Sigma Aldrich, 15µM okadaic acid OA - Enzo Life Sciences or Sigma, 63 µg/ml NaF and 315 µg/ml NaV). 10µl per condition of p13-sepharose beads^59,45,46^ or glutathione-agarose beads (G4510, Sigma), used as control, were pre-washed and incubated with either prophase oocyte extract or PBS (13.7 mM NaCl, 2.7 mM KCl, 4.3 mM KH_2_PO_4_, 1.4 mM Na_2_HPO_4_, pH 7.4) at 4°C for 18 hrs under rotation. Beads were washed under rotation three times with buffer E (250 mM NaCl, 0.2% Triton, Antiproteases cOmplete^TM^ Sigma Aldrich, 10 µM OA, 42 µg/ml NaF and 210 µg/ml NaV) for 5 min at 4°C and three times with EB. All the samples were incubated at 30°C for 30 min and eluted.

### In extract kinase assay

40 prophase oocytes were lysed at a ratio of one oocyte in 5µl of buffer F (80 mM β-glycerophosphate pH 7.3, 20 mM EGTA, 15 mM MgCl_2_, anti-proteases - cOmplete^TM^ Sigma Aldrich, 10 µM okadaic acid OA - Enzo Life Sciences or Sigma, 42 µg/ml NaF and 210 µg/ml Na_3_VO_4_) supplemented with 1 mM ATP. Half of the lysate was incubated in the presence of 40 µg of PKI for 15 min at 18°C. Δ126-Cyclin B2 or Arpp19 was then added at 100 ng per oocyte equivalent, and the reactions were incubated for 60 min at 18°C. Samples corresponding to one oocyte equivalent were analyzed by western blot.

### *In vitro* kinase assays

Unless otherwise indicated, *in vitro* kinase assays were performed in a reaction buffer G (20 mM HEPES, 20 mM MgCl₂, 1 mM ATP, 2 mM β-mercaptoethanol, pH 7.4) and reactions were incubated for 30 min at 30°C. To assess PKA-mediated phosphorylation of Cyclin B2, Cyclin B1 and Cyclin A2, PKAc (Promega, V5161) (50 ng) was incubated with 50 ng of either Δ126- Cyclin B2 or Δ126-S271A Cyclin B2, 100 ng of either commercial Cdk1-Cyclin B1 (Abcam, Ab271456) or Cdk1-Cyclin A2 (Ab64299). As indicated, 1 µg of V5-PKI or 1.5 µg of V5- Cip1 was added to the reactions. For identification of the PKA phosphorylation site on Cyclin B2 by mass spectrometry, kinase assays were performed under the same conditions using increased amounts of enzyme and substrate: 500 ng of PKAc and 500 ng of Δ126-Cyclin B2. The effect of PKA phosphorylation on the ability of Cdk1-Cyclin B1 to phosphorylate GST- PP1 was assessed as previously described in^29^, using 20 ng of Cdk1/Cyclin B1 phosphorylated as described above. To validate the phospho-specific antibody against Cyclin B2 phosphorylated at S271, increasing amounts of Δ126-Cyclin B2 (12.5, 25, 50, 100 or 200 ng) were incubated in the presence of 50 ng of PKAc under the same reaction conditions. To assess the effect of Cyclin B1 phosphorylation by PKA on the ability of Cdk1 to induce multiple phosphorylation of Cdc25C, Cdk1-Cyclin B (30 ng) was first incubated in the presence or in the absence of 15 ng of PKAc for 30 min at 25°C to enable PKA-mediated phosphorylation of Cdk1-Cyclin B. Recombinant PKI (100 ng) was subsequently added to inhibit PKAc activity. PKA-phosphorylated or non-phosphorylated Cdk1-Cyclin B was then incubated under the same reaction conditions in the presence of 125 ng of recombinant Cdc25C (residues 89–342), previously phosphorylated by PKAc. Samples were collected after 5, 15, 30 and 60 min incubation for subsequent analysis. The center of mass was calculated using the custom script *Central Mass Calculator.py*. The data were interpolated using a one-phase association equation and the half-time was determined using Prism10.

### Gel filtration

Oocytes were injected or not with Δ90-Cyclin B2, Δ90-S271A-Cyclin B2 or Δ90-S271D- Cyclin B2. 50 prophase-arrested oocytes were homogenized in 200 μl of buffer H (80 mM β - glycerophosphate, 30 mM NaCl, 10 mM EDTA, 1 mM DTT, pH 7.4) supplemented with anti-proteases - cOmplete^TM^ Sigma Aldrich, 3 mM DTT, 42 µg/ml NaF and 210 µg/ml Na_3_VO_4_. 80% of the extract was recovered after 20.000 g centrifugation for 10 min at 4°C and further ultracentrifuged in a TLA100 rotor at 53.000 rpm for 15 min at 4°C. 120 μl of extract were injected into a Superose 12 gel filtration column. 20 ml were eluted and collected in 500 μl fractions. Proteins were precipitated by deoxycholate at 0.02% final concentration and TCA at 10%. After 1 hr incubation at 4°C, fractions were centrifuged at 15.000 g for 20 min at 4°C. Pellets were washed in acetone and resuspended in 40 µl EB. Samples were analysed by western blot.

### Data availability

Data are available via ProteomeXchange with identifier PXD081389

## ACKNOWLEDGMENTS

This work was supported by the National Centre for Scientific Research (CNRS) and Sorbonne University, the ARC foundation grant ARCPJA2023080006901 to EMD, a 4^th^ year PhD fellowship from ARC foundation to MS, the National Research Agency (ANR) (ANR-23-CE12-0045-01) to EMD.

## AUTHOR CONTRIBUTIONS

**Martina Santoni:** Methodology; Experiments; Formal Analysis; Data Curation; Writing; **Tran Le:** Experiments; Formal Analysis; **Catherine Jessus:** Writing; Supervision; **Veronique Legros**: Mass spectrometry analysis, Writing; **Guillaume Chevreux:** Mass spectrometry analysis, Writing; **Enrico Maria Daldello**: Conceptualization; Methodology; Formal Analysis; Data Curation; Writing; Supervision.

**Sup. Fig. 1: Optimization of Turbo-ID approach.**

(A) Prophase-arrested oocytes were injected with mRNA encoding Turbo or Turbo-PKAc. After 18 hrs, oocytes were stimulated with progesterone and NEBD was scored by monitoring the appearance of the white spot. (B) Prophase-arrested oocytes were injected with mRNA encoding Turbo or Turbo-PKAc. After 18 hrs oocytes were incubated with exogenous biotin for the indicated times. Biotinylated proteins were recovered by streptavidin pull-down and identified by LC–MS/MS. The volcano plots display log_2_ fold-change (Turbo-PKAc vs Turbo) on the x-axis versus the relative abundance of the protein expressed as the log_10_ of the sum of the areas of the peptide of each protein in the y-axis. Known PKA interactors: green; Cyclin B2: red; carboxylases: blue. (C) The average position of total proteins (grey), of carboxylases (blue), and of PKA known interactors (green) were plotted for each time point in the x-y space described in panel B. (D) Prophase-arrested oocytes were injected with mRNA encoding Turbo or Turbo-PKAc (T-PKAc). After 18 hrs, oocytes were incubated in the presence of exogenous biotin for 60 min. Biotinylated proteins were recovered by streptavidin pull-down. Lysates (INPUT) and supernatant after pull-down (SPN) were immunoblotted with anti-HA (detecting Turbo and Turbo-PKAc), anti-PKAc (detecting Turbo-PKAc and endogenous PKAc) and anti-RRxS/T (PKA substrate motif) antibodies, and probed with streptavidin–HRP to visualize biotinylated proteins.

**Sup. Fig. 2: Cyclin Bs, but not Cyclin A, are phosphorylated by PKA**

(A) Different quantities of V5-Δ126-Cyclin B2 were incubated for 30 min at 30°C with PKAc. Kinase assay reaction was immunoblotted with anti-pS271-Cyclin B2, anti-Cyclin B2 and anti-PKAc antibodies. (B) Quantification of endogenous levels of Cyclin B2 in prophase-arrested oocytes. Prophase-arrested oocyte extracts were compared with different concentration of V5-Δ126-Cyclin B2 recombinant protein. (C) Prophase extracts were supplemented or not with PKI. After a 15 min incubation, 100 ng of recombinant V5-Arpp19 was added for 60 min. Lysates were immunoblotted with anti-pS109-Arpp19, anti-V5, anti-PKAc and anti-karyopherin antibodies. (D) Alignment of *Xenopus* and human Cyclin B and Cyclin A sequences. The PKA consensus motif and the PBP are indicated in red and black respectively. (E) PKAc was preincubated or not in the presence of PKI for 30 min. Human GST-Cyclin B1-Cdk1 and GST-Cyclin A2-Cdk1 complexes were preincubated or not with the Cdk1 inhibitor Cip1 for 30 min. Both samples were mixed and further incubated for 30 min. Western blots were performed with anti-RRxpS/T, anti-PKAc, anti-GST (to detect GST-Cdk1, GST-Cyclin B1, GST-Cyclin A, GST-Cip1), and anti-V5 (to detect V5-PKI) antibodies.

